# A novel domain within the CIL regulates egress of IFITM3 from the Golgi and prevents its deleterious accumulation in this apparatus

**DOI:** 10.1101/2021.03.29.437470

**Authors:** Li Zhong, Rustem Uzbekov, Chloé Journo, Philippe Roingeard, Andrea Cimarelli

**Author notes:** Correspondent footnote: correspondence should be addressed to Andrea Cimarelli, CIRI, 46 Allée d’Italie, 69364 Lyon, France.

## Abstract

The InterFeron-Induced TransMembrane proteins (IFITMs) are broad viral inhibitors that protect cells by preventing viral-to-cellular membrane fusion and they belong to the dispanin/CD225 family that includes vesicle trafficking regulators and proteins of unknown functions into four subfamilies (A-D). In this study, we uncover a novel domain that regulates the egress of IFITM3 from the Golgi and that is required to prevent IFITM3-driven v- to t-SNAREs membrane fusion inhibition and Golgi dysfunctions.

The S-x-K-x-R-D domain is conserved among vertebrate members of the dispanin/CD225 A subfamily that regroups all IFITMs and through the study of mutations identified in patients affected by paroxysmal kinesigenic dyskinesia (PKD), we determine that it is functionally conserved also in PRRT2, member of the B subfamily.

Overall, our study defines a novel domain that regulates the egress of dispanin/CD225 members from the Golgi and stresses the importance that regulation of this process bears to preserve the functions of this apparatus.

## Introduction

The InterFeron-Induced TransMembrane proteins (IFITMs) are broad viral inhibitors that prevent membrane fusion between viral and cellular membranes, thus protecting the cell from infection ^1^. IFITMs belong to the dispanin/CD225 family of proteins that originated from metazoan lineages and diversified into four distinct subfamilies (A through D) that include proteins of known and yet unknown functions ^2^. The A subfamily regroups all IFITMs which in humans are: IFITM1, 2 and 3, that we will refer to hereafter as IFITMs, that are interferon (IFN)-induced proteins essentially studied in the context of viral infection ^3^; IFITM5 that plays undefined roles in Osteogenesis imperfecta type V, a bone-specific disease^4^ and IFITM10 whose functions are unknown. An additional known member of the dispanin/CD225 family is the neuron specific PRoline-Rich Transmembrane protein 2 (PRRT2, B subfamily) that is involved in neurotransmitter vesicles regulation and has been genetically linked to benign familial infantile seizures (BFIS) and to paroxysmal kinesigenic dyskinesia (PKD), the most common type of paroxysmal movement disorder ^5^.

Members of the dispanin/CD225 family are characterized by a similar structure consisting of an intramembrane domain (IMD, previously defined as transmembrane domain 1, TM1), a cytoplasmic intracellular loop (CIL) and a transmembrane domain TMD (previously defined as transmembrane domain 2, TM2, as schematically shown in Fig. 1a for IFITM3). By virtue of this structure, IFITMs decorate cellular membranes in which they act as membrane fusion inhibitors. The distribution and activities of IFITMs on cellular membranes are highly regulated and are driven not only by distinct N and C termini responsible for the more prominent distribution of individual IFITMs at the plasma or at inner membranes as is the case for IFITM1 with respect to IFITM2 and 3, respectively, but also by several post-translational modifications and in particular palmitoylation on cysteine residues, ubiquitination and methylation on lysines, as well as phosphorylation on a specific tyrosine residue ^6–10^.

**Fig. 1.**
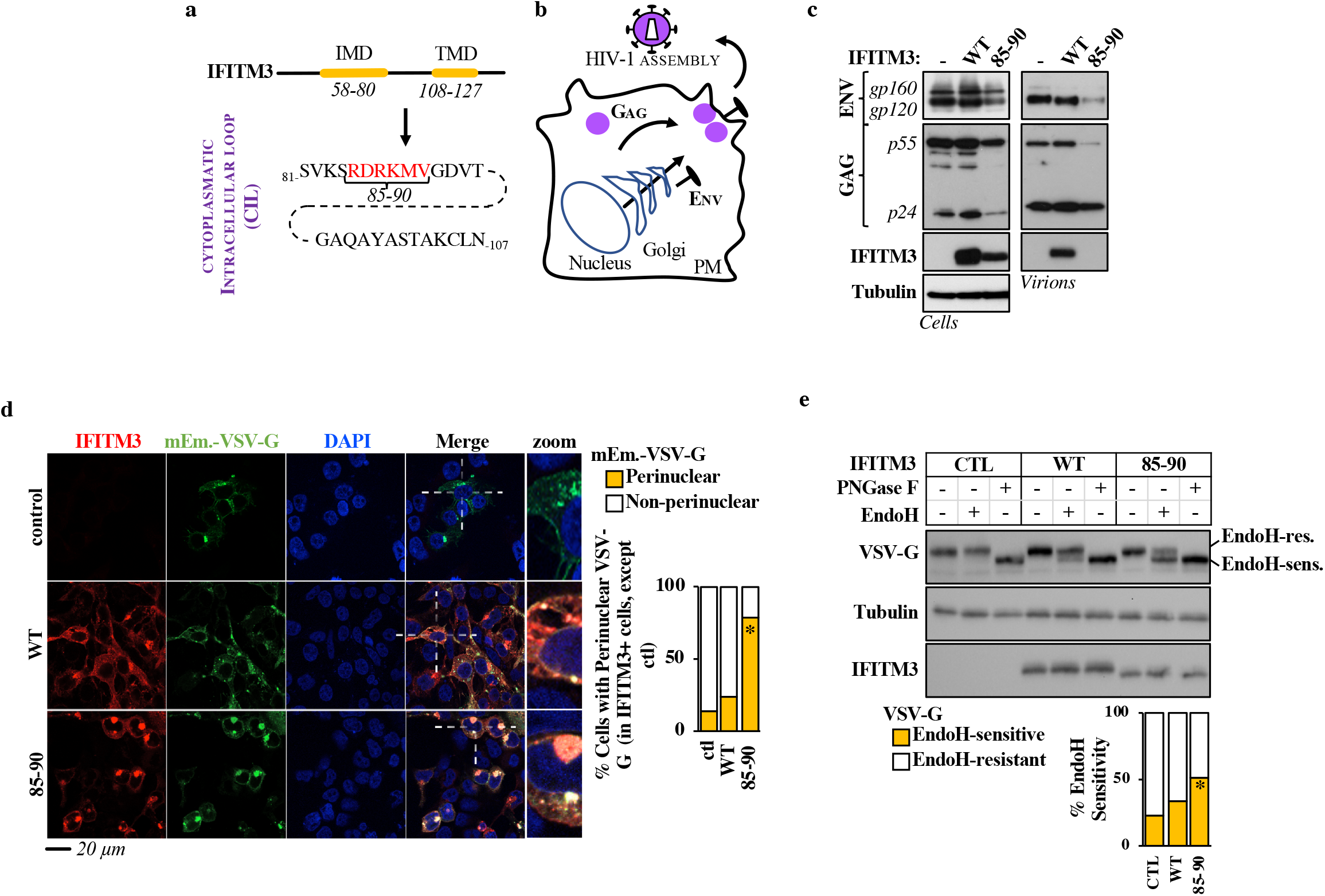
A novel gain-of-function IFITM3 mutant induces secretory pathway defects that affect glycoproteins trafficking. **a)** Genomic structure of IFITM3 with a highlight on amino acids of the cytoplasmic intracellular loop (CIL). In mutant 85-90, red residues have been mutated to alanine. **b)** Schematic representation of HIV-1 virion assembly. **c)** HEK293T cells were transfected with DNAs coding for the HIV-1 proviral clone NL4-3 along with control, WT or 85-90 mutant IFITM3. Cells were lysed 48 hours after transfection, while virion particles released in the supernatant were first purified by ultracentrifugation through a 25% sucrose cushion. Both cellular and viral lysates were analyzed by WB. **d)** HEK293T cells were ectopically transfected with DNAs coding for the indicated IFITM3s along with the G protein of the Vesicular Stomatitis virus fused to the mEmerald fluorescent reporter (mEm.-VSV-G), prior to confocal microscopy analysis twenty-four hours later. Representative confocal microscopy images and graph presenting the proportion of double-positive cells displaying perinuclear accumulation of mEm.-VSV-G (binary scoring; three independent experiments; between 50 and 100 cells scored per sample). *, p value <0.0001 following a one-way Anova, Tukey’s multiple comparison test; non-statistically significant differences, not shown. **e)** Lysates obtained from cells expressing VSV-G along with control, WT and mutant IFITM3 were either untreated or treated with EndoH or PNGaseF prior to WB and densitometry quantification of the EndoH-sensitive and -resistant VSV-G forms. Panels present representative results, while the graph presents the proportions of EndoH-sensitive and -resistant VSV-G obtained in 5 independent experiments. *, p value of 0.0021 following a one-way Anova, Tukey’s multiple comparison test over control. Uncropped blots and source data are provided in the relevant section.

More recently, IFITMs have been linked to cellular functions independent from their protective role against viral infection in the regulation of glucose metabolism in mice, of phosphatidylinositol 3-kinase (PI3K) mediated signaling in B cells ^11^ and of trophoblast fusion during placental formation *in vivo* ^12^, overall supporting the notion that IFITM proteins likely act well beyond a viral context.

In this study, through an in-depth characterization of a novel gain-of-function IFITM3 mutant, we have been able to identify a novel protein domain with the CIL of IFITM3 that regulates its egress from the Golgi apparatus. Mutations within this domain lead to the retention of IFITM3 in this organelle, where IFITM3 interferes with the fusion mediated between vesicle and target Soluble N-éthylmaleimide-sensitive-factor Attachment protein REceptor (v- and t-SNAREs), impairing the functionalities of the Golgi secretory pathway.

Functional as well as in silico analyses allowed us to precisely define the Golgi egress _81_S-x-K-x-R-D_86_ domain which is positioned after the IMD, at the beginning of the CIL and it is conserved across vertebrate members of the dispanin/CD225 A subfamily. Through the analysis of mutations identified in patients affected by PKD, we do show here that, despite no conservation at the amino acid level, this domain is functionally conserved also in PRRT2, a member of the B subfamily, thus raising the possibility of a general functional role in Golgi trafficking common to dispanin/CD225 subfamilies.

Overall, these results identify a domain that regulates egress of IFITM and IFITM-like proteins from the Golgi and given the deleterious effects that dysregulation of this process bears on the overall functionalities of this apparatus, they highlight the importance that correct regulation of IFITM trafficking has for the cell.

## Results

### A gain of function IFITM3 mutant affects glycoprotein trafficking

During a previous mutagenesis study of the broad viral inhibitor IFITM3, we identified a mutant in the cytoplasmic intracellular loop region of IFITM3 (CIL, mutant 85-90 in which six consecutive residues were mutated to alanine, Fig. 1a) that exerted a strong inhibitory effect on the incorporation of HIV-1 Envelope glycoproteins at the surface of released virion particles^13^. At its simplest, the formation of infectious HIV-1 virion particles requires the gathering on the plasma membrane of two classes of structural viral proteins: Gag (and Gag-Pro-Pol, not marked here for clarity’s sake) and Env (Fig. 1b). While Gag is translated from cytoplasmic mRNAs from free ribosomes, the Envelope glycoprotein undergoes co-translational ER translocation and reaches the plasma membrane following the ER-Golgi secretory pathway14. As such, loss of Envelope incorporation in virion particles may reflect potential effects of IFITM3 along this axis that we decided to investigate further.

Indeed, when expressed in HEK293T cells undergoing HIV-1 virion assembly (Fig. 1b-c), the 85-90 IFITM3 mutant induced only a moderate, yet detectable, decrease in the levels of cell-associated Gag and Env proteins when compared to control or WT IFITM3-expressing cells, but as expected it induced a drastic decrease in the amount of Env glycoprotein incorporated in purified virion particles (Fig. 1c). From this starting observation, we decided to determine whether this defect could be more generally reflective of an interference with Golgi-mediated trafficking, independently from HIV-1. To this end, the 85-90 IFITM3 mutant was co-expressed along with a fluorescent G protein of the Vesicular Stomatitis Virus (mEm.-VSV-G, Fig. 1d), prior to confocal microscopy analysis. Under these conditions, a strong perinuclear accumulation of mEm.-VSV-G was observed in cells expressing the mutant IFITM3, overall suggesting a defect in Golgi-mediated trafficking (Fig. 1d).

To further support this result, we analyzed the EndoH susceptibility of a non-tagged VSV-G followed by WB analysis, as an independent and commonly used technique to appreciate the progression of glycosylated proteins from the ER through the Golgi (Fig. 1e). Under these conditions, expression of WT IFITM3 led to a small but detectable accumulation of EndoH-sensitive VSV-G when compared to control cells which however did not reach statistical significance under the conditions used here. However, the proportion of EndoH-sensitive VSV-G protein was significantly increased upon expression of the 85-90 IFITM3 mutant, indicating a strong defect in the ER-Golgi secretory pathway in the presence of this mutant protein.

Overall, these results indicate that the expression of the 85-90 IFITM3 mutant affects the normal functionalities of the cellular ER-Golgi secretory pathway, highlighting it as a novel gain-of-function IFITM3 mutant.

### The 85-90 IFITM3 mutant accumulates in and leads to structural changes of the Golgi apparatus

The 85-90 IFITM3 mutant did not significantly colocalize with the ER, nor with endosomal vesicles (Extended data Fig. 1). Instead, this mutant was strongly localized at the cis-Golgi (GM130 marker, Fig. 2a and 2b for a 3D-reconstruction of positive cells) and induced gross morphological changes in the appearance of the Golgi: punctiform in control and WT-IFITM3 expressing cells and inflated in cells expressing the 85-90 IFITM3 mutant. These structural changes were not only apparent after 3D-reconstruction of positive cells following confocal microscopy analysis (Fig. 2b), but also after electron microscopy analysis which more precisely indicated that the shape of the Golgi apparatus had lost its classical punctiform distribution in favor of an expansion of large vesicles (Fig. 2c).

**Fig. 2.**
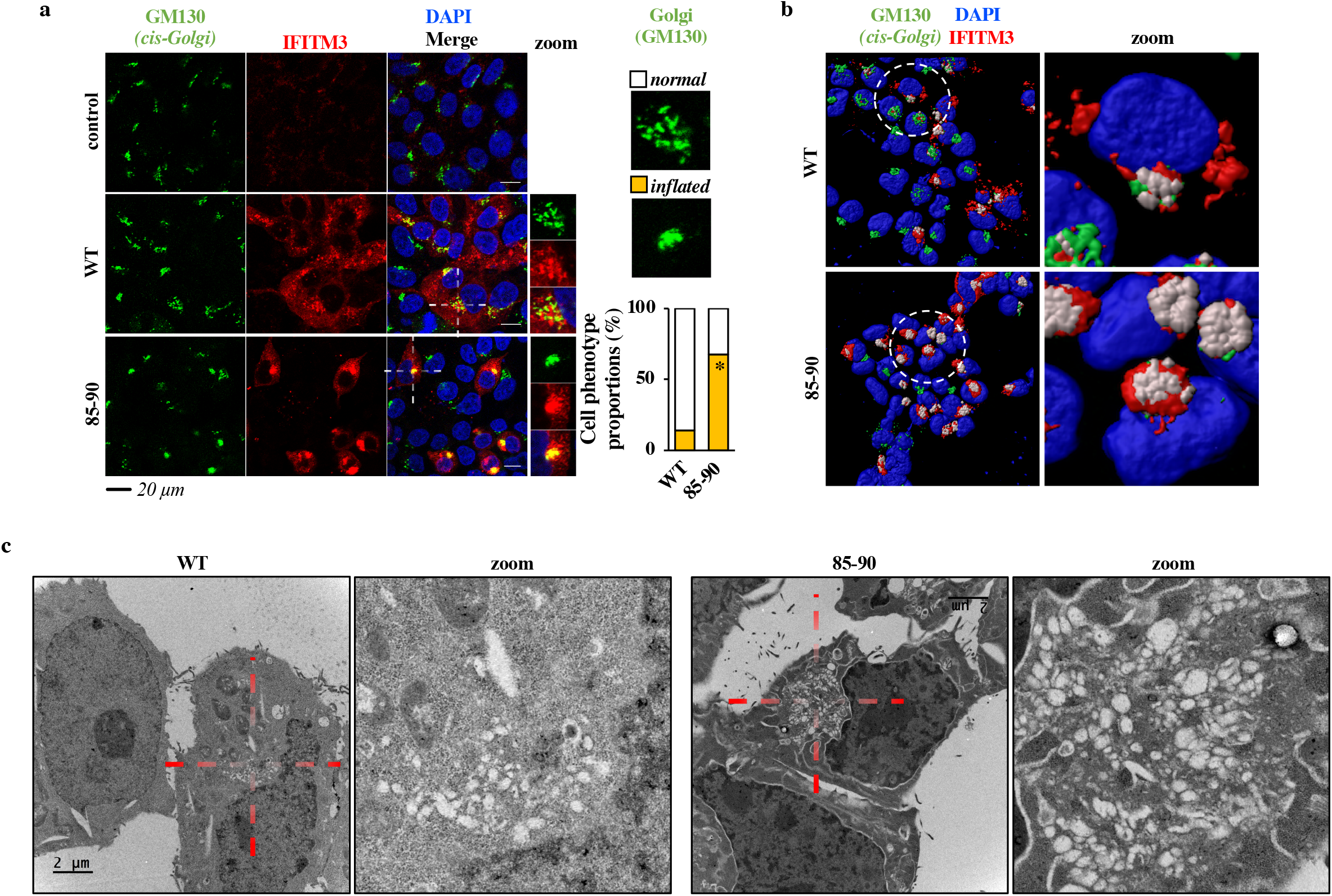
The 85-90 IFITM3 mutant colocalizes with and induces gross morphological changes in the Golgi. **a and b)** Confocal microscopy analysis and 3D-reconstruction of cells expressing the indicated IFITM3 proteins. Representative images and graph presenting phenotype proportions in at least 100 cells scored per condition (binary scoring: inflated or normal Golgi, three independent experiments). *, p=8.3×10-9 following an unpaired, two-tailed Student t test. **c)** Representative electron microscopy analyses of HEK293T cells expressing WT or 85-90 IFITM3 proteins. The region enlarged at the right of each panel corresponds to the red inset. Source data is provided in the relevant section.

### The 85-90 IFITM3 mutant affects fusion between t and v-SNAREs

One of the key features of IFITM family members is their ability to interfere with viral-to cellular membranes fusion^15–17^. We thus hypothesized that Golgi dysfunctions could be caused by the interference of IFITM3 with the ability of v-SNARE vesicles to fuse to t-SNAREs compartments, a process that is key to maintain protein and membrane fluxes, that is key for the integrity of the Golgi and that is reminiscent of the fusion between membranes that is observed after the engagement of a viral envelope glycoprotein with its cellular receptor^18,19^. To test this hypothesis, we analyzed the ability of the 85-90 IFITM3 mutant to inhibit the fluorescent resonance energy transfer (FRET) that occurs upon fusion between GS15-CFP and ERS24-YFP bearing vesicles (v-and t-SNAREs, respectively), as a measure of membrane fusion inhibition in this compartment, as described ^20^ (Fig. 3a). Given that this analysis was carried out in living cells, an excess of the 85-90 IFITM3 mutant was used to ensure that the majority of GS15-CFP and ERS24-YFP double positive cells were also IFITM3-positive. Under these conditions, a statistically significant decrease in FRET was observed upon expression of the 85-90 IFITM3 mutant with respect to control cells, indicating that this mutant does indeed interfere with v-to t-SNAREs fusion, in agreement with the major function ascribed to IFITM3 as a membrane fusion inhibitor.

**Fig. 3.**
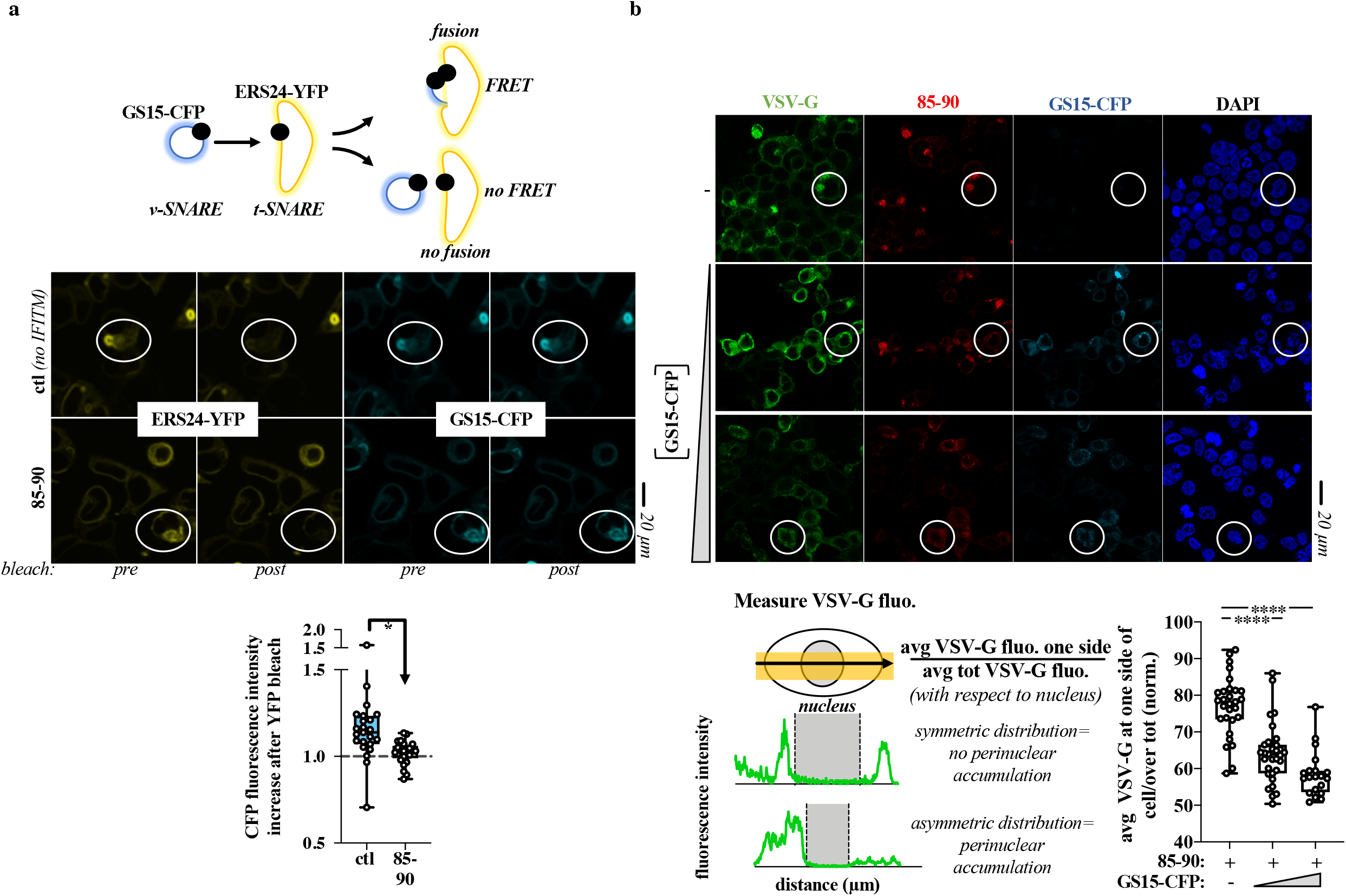
The 85-90 IFITM3 mutant impairs fusion between v-and t-SNAREs. **a)** Schematic representation of the FRET experiment and representative images pre and post ERS24-YFP bleach. FRET experiments were performed on live HEK293T cells in which an excess of DNA coding for the 85-90 IFITM3 mutant was used over GS15-CFP and ERS24-YFP (five-fold excess) to ensure that the majority of cells analyzed were also IFITM3-positive. The boxes and whiskers plot presents the normalized decrease and values distribution in FRET observed in control and 85-90 IFITM3 expressing cells from three independent experiments and >20 cells examined. *, p value of 0.0008, following an unpaired two-tailed Student t test. **b)** HEK293T cells expressing constant amounts of 85-90 IFITM3 and VSV-G were co-transfected with increasing amounts of GS15-CFP coding DNA, prior to confocal microscopy analysis (GS15/85-90 ratios of 0.2 and 0.5). Given that VSV-G accumulation in the Golgi leads to its asymmetrical perinuclear distribution in the cell, VSV-G fluorescence was determined for each cell according to the presented scheme. The fluorescence measured over distance in the cell was used to calculate the proportion of VSV-G protein present at one side (the left side was set as the side with higher protein accumulation). Representative pictures and graph presenting the distribution of VSV-G at one side of the cell in triple-positive cells (n=2 with 20 to 31 cells analyzed per condition). White circles highlight example cells of increased FRET (a) and of phenotypic differences in the perinuclear accumulation of VSV-G (b), respectively. ****, p value of <0.0001 following a one-way Anova, Tukey’s multiple comparison test over 85-90 IFITM3 condition with no GS15-CFP. Source data is provided in the relevant section.

To further support this observation, we overexpressed the v-SNARE GS15 and determined whether this could lead to Golgi trafficking normalization despite the presence of the 85-90 IFITM3 mutant. Indeed, v-SNAREs overexpression is known to bypass membrane fusion defects between Golgi membranes ^21–23^ and we thus hypothesized it could relieve the defect imposed by the IFITM3 mutant. To this end, increasing levels of GS15 were expressed along with a constant amount of both 85-90 IFITM3 mutant and VSV-G, prior to the analysis of the distribution of VSV-G within triple-positive cells (Fig. 3b). Under these conditions, GS15 was able to relieve the perinuclear accumulation of the VSV-G glycoprotein induced by the IFITM3 mutant and was also able to act similarly on the 85-90 IFITM3 mutant itself, supporting our contention that a main disruptive event due to the accumulation of the 85-90 IFITM3 mutant in the Golgi is the inhibition of v-to t-SNAREs fusion.

### The 85-90 mutation in the IFITM3 CIL defines a domain that regulates IFITM3 egress from the Golgi

To assess whether the 85-90 mutation induced retention or relocalization of IFITM3 at the Golgi, we performed a time-course analysis of both WT and mutant IFITM3 from early to late time points after ectopic DNA transfection (Fig. 4). While both proteins exhibited a predominant Golgi distribution at 6 hours post transfection, such concentration gradually diminished in the case of WT IFITM3, in agreement with its functional exit from the Golgi. On the contrary, little changes were observed over time in the case of the 85-90 IFITM3 mutant that remained in this compartment, supporting the notion that the 85-90 mutation induces the retention, rather than the redistribution, of IFITM3 in the Golgi.

**Fig. 4.**
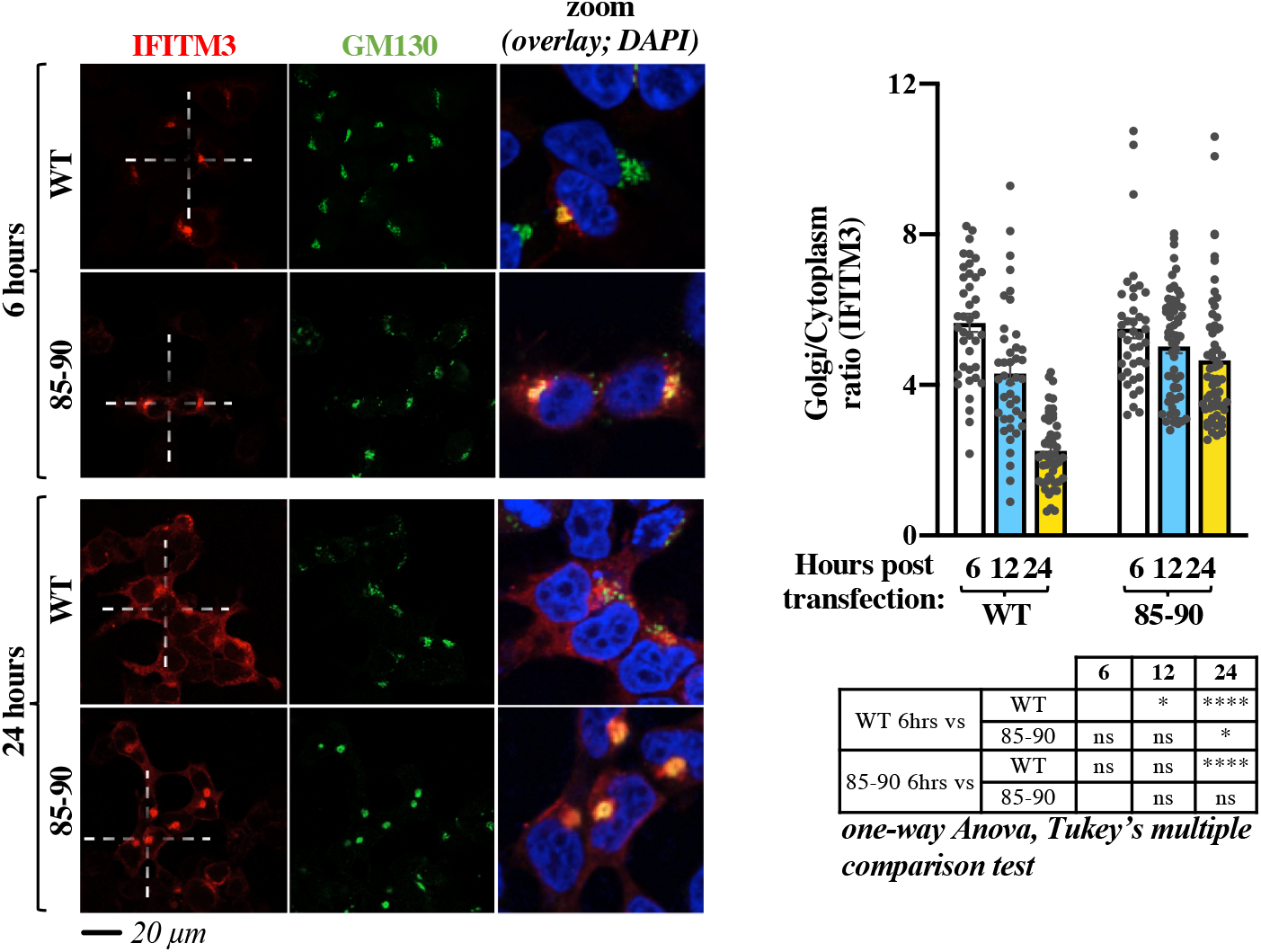
Time course confocal microscopy analysis indicates that the 85-90 IFITM3 mutation affects the normal egress of IFITM3 from the Golgi. HEK293T cells were examined by confocal microscopy at different times post ectopic DNA transfection to determine the localization of IFITM3 proteins over time. Representative pictures and graph presenting the distribution of IFITM3 in the Golgi (calculated as a Golgi/cytoplasm ratio on 39 to 62 cells per time point and per condition in two independent experiments, AVG, SEM and individual values). The table presents p values obtained after a one-way Anova, Tukey’s multiple comparison test between the indicated conditions: ns: non significant; *, p value <0.05; ****, p value <0.0001. Source data is provided in the relevant section.

In agreement with this observation, incubation of cells with the broad trafficking inhibitor Monensin induced accrued Golgi accumulation of WT IFITM3, but as expected did not modify the distribution pattern of the 85-90 mutant (Extended data Fig. 2).

Overall, these results concur in indicating that the 85-90 mutation defines a protein domain within the CIL that regulates the normal egress of IFITM3 from the *cis*-Golgi.

### Fine mapping of the Golgi egress domain of IFITM3 within the CIL

To more precisely map the residues that regulated the egress of IFITM3 from the Golgi, individual amino acids spanning the entire CIL were mutated to alanine and mutants were analyzed by confocal microscopy (Fig. 5a-b and Extended data Fig. 3, alanine residues in the CIL were instead mutated to glycines). With the exception of two mutants that were barely detectable by WB (D92A and G95A), the remaining mutants were expressed to detectable levels upon WB analysis. We noticed that the D92A mutant acquired also a nuclear localization unusual for IFITM proteins, but this was not characterized further. This mutagenesis study identified a number of residues whose mutation led to accrued accumulation of IFITM3 into the Golgi (S81, V82, K83, S84, R85, D86, A96, Q97, Q98, S101, T102, A103, K104, L106 and N107), indicating that several residues within the CIL participate to this phenotype. Interestingly however, the mutations that resulted in levels of IFITM3 localization in the Golgi equivalent to those of the 85-90 mutant were essentially clustered in a patch (from residues S81 to D86) that was posited at the boundary between the end of the intramembrane domain of IFITM3 (IMD) and the start of the CIL and that overlapped with the 85-90 stretch.

**Fig. 5.**
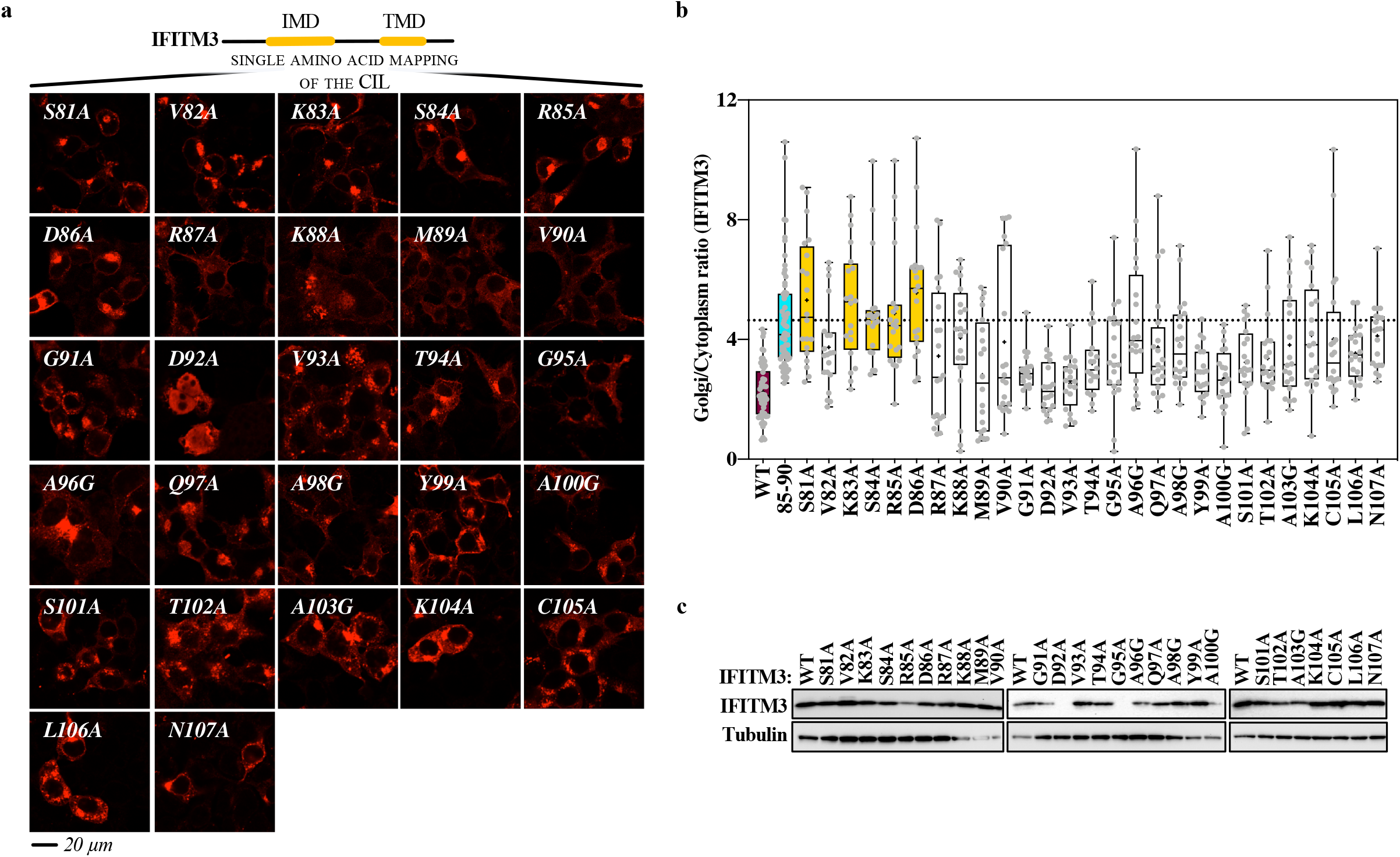
Mutagenesis of the entire CIL of IFITM3 finely maps the domain involved in the egress of IFITM3 from the Golgi. **a)** Individual amino acid of the CIL were changed to alanine or to glycine when alanine residues were present, prior to confocal microscopy analysis. Representative pictures are shown here (IFITM3 only) while their co-localization with the cis-Golgi marker GM130 is shown in the extended data Figure 3. **b)** Quantification of the IFITM3 proportion in Golgi is provided as a Golgi/cytoplasm ratio in the box and whisker plot (20 cells per mutant in two to three independent experiments analyzed). Asterisks and lines within the box indicate averages and median values, respectively. Yellow boxes indicate mutants with Golgi/cytoplasm IFITM3 ratios equivalent to the 85-90 IFITM3 mutant and non-statistically significant when compared to 85-90 following an ordinary one-way Anova, Dunnett’s multiple comparison test. **c)** Representative WB analysis of mutant proteins. Uncropped blots and source data are provided in the relevant section.

Thus, the complete mutagenesis of the CIL allowed us to more precisely refine the domain responsible for the exit of IFITM3 from the Golgi (_81_SxKSRD_86_).

### A generalized Golgi egress domain is conserved among members of the dispanin/CD225 subfamily A and is functionally conserved in PRRT2, a member of the dispanin/CD225 subfamily B, genetically associated to paroxysmal kinesigenic dyskinesia (PKD)

IFITMs belong to the larger dispanin/CD225 family which is itself divided into four subfamilies (A to D, Fig. 6a) ^2^. The A subfamily is composed of IFITM1, 2 and 3, as well as IFITM5 and IFITM10, which are not IFN regulated and exert unclear functions that in the case of IFITM5 are associated to Osteogenesis imperfecta type V, a bone-related genetic disease^4^. The alignment of residues spanning the IFITM3 SxKSRD domain in human IFITMs, as well as their comparison in vertebrates (Fig. 6a, aligned sequences on the left for human IFITMs and Logo on the right for vertebrate IFITMs, respectively) indicated the presence of a consensus sequence conserved across vertebrate members of the dispanin/CD225 A subfamily: S-x-K-x-R-D. This patch was not conserved at the amino acidic level in the remaining subfamilies.

**Fig. 6.**
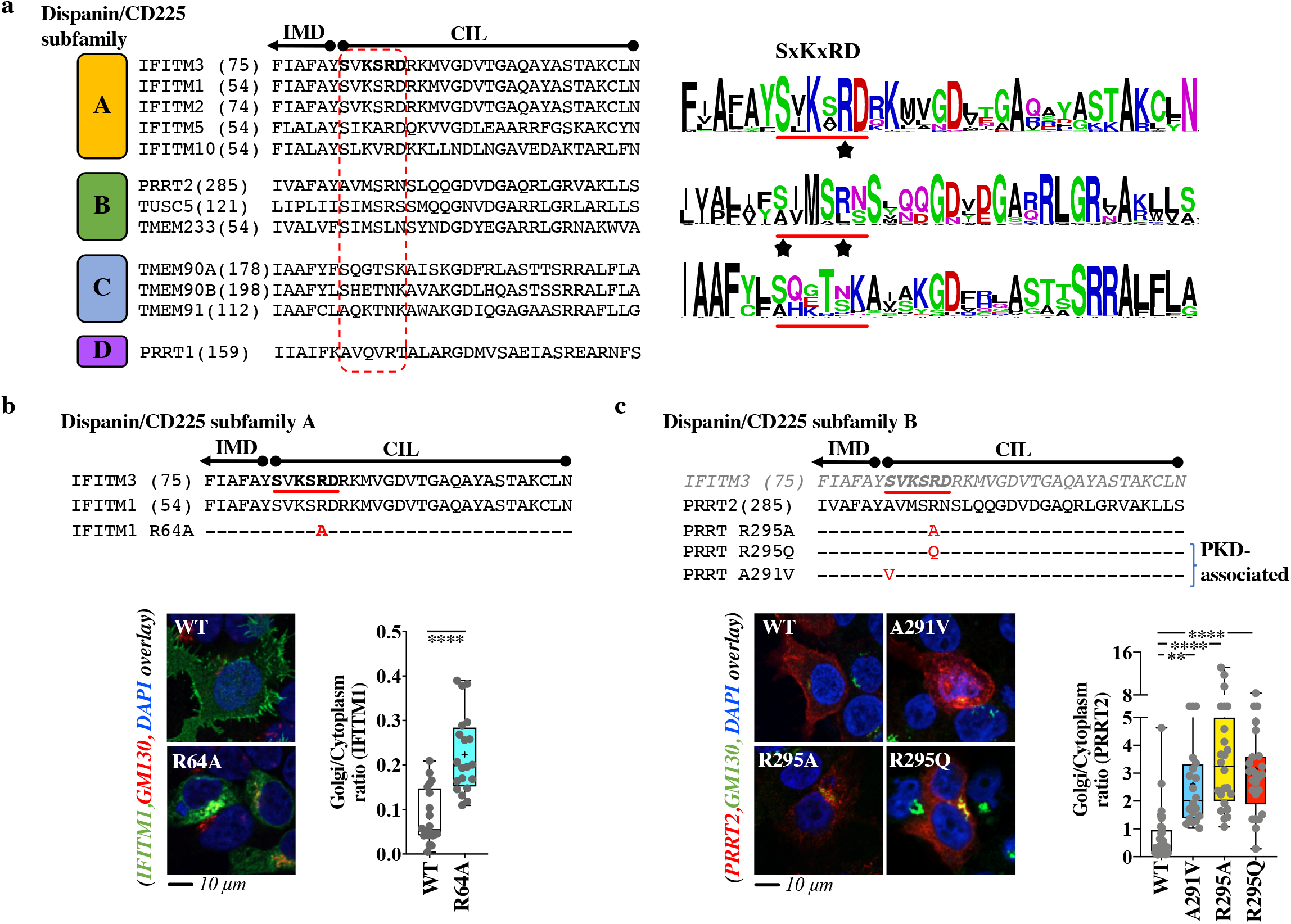
The Golgi egress domain of IFITM3 is conserved across vertebrate members of the dispanin/CD225 subfamily A and is functionally conserved in PRRT2, a member of the B subfamily as shown by genetic mutations associated to PKD. **a)** Alignment of the indicated portions of human members of the different dispanin/CD225 subfamilies. The relevant domain is circled in red. The position of the first amino acid of each sequence is shown within parentheses. Left, logo files obtained after alignment of vertebrate orthologues of the indicated genes for each subfamily, except D. **b)** Single point mutation and confocal microscopy analysis of the Golgi egress domain of IFITM1. **c)** Point mutations and confocal microscopy analysis of PRRT2 mutants in the region corresponding to the Golgi egress domain of IFITM3. R295Q and A291V are genetic mutations associated to PKD. Representative pictures are shown for each protein, while separated channels are provided in the extended data Fig. 4. Whiskers and boxes plots of the quantification of the proportion of each protein in the Golgi as a Golgi/cytoplasm ratio (25-75 percentiles with individual cells represented as dots; group average indicated by an asterisk, n=20 in three independent experiments). Colored boxes represent statistical significant differences after a two-tailed Student t test or a One-way Anova Dunnett’s multiple comparison test (b and c, respectively) of the examined mutant over WT. **, p value of 0.0019; ****, p value <0.0001. Source data is provided in the relevant section.

To determine whether this consensus domain exerted similar functions in other IFITM proteins belonging to the A subfamily, a single point mutation was introduced in the corresponding domain of IFITM1 (R64A), that contrarily to IFITM3 is essentially localized at the plasma membrane due to the lack of specific trafficking domains at its N-terminus when compared to IFITM3. Under these conditions, the introduction of a single point mutation in the IFITM Golgi egress domain was sufficient to drive the redistribution of IFITM1 from the plasma membrane to the Golgi, as observed by confocal microscopy analysis (Fig. 6b).

Thus, *in silico* and functional analyses indicate the presence of a conserved S-x-K-x-R-D domain important for the exit of members of the dispanin/CD225 A subfamily from the Golgi.

Despite the fact that the functions of certain proteins belonging to the dispanin/CD225 family remain unknown (as is the case for TMEME90A, TMEM91, TMEM233 and TMEM265), the information existing on other members (TUSC5, TMEM90B and PRRT2) indicate functions revolving around vesicular trafficking and vesicular behavior ^5 24,25^. Among them, PRRT2 acts as a regulator of neurotransmitter vesicles in neuron cells and is genetically linked to paroxysmal kinesigenic dyskinesia (PKD, the most common type of paroxysmal movement disorder), as well as benign familial infantile seizures (BFIS). While the Golgi egress domain identified in IFITM proteins did not appear to be conserved at the amino acid level in other subfamilies, we decided to investigate whether it could be functionally conserved in PRRT2 as a prototype member of the B subfamily. Indeed, we noticed that two mutations in PRRT2 have been identified in patients affected by PKD in the same CIL region at the boundary between the IMD and the CIL (A291V and R295Q, corresponding to the underlined positions 1 and 5 of the <u>S</u>-x-K-x-<u>R</u>-D domain) ^26,27^.

To test this hypothesis, the R295Q and 1291V mutations were introduced along with a R295A alanine mutation in PRRT2 and analyzed by confocal microscopy (Fig 6c). While WT PRRT2 exhibited a predominant plasma membrane distribution, introduction of the above-mentioned mutations led to accrued redistribution of PRRT2 in the Golgi, similarly to what described for IFITM3.

Thus, these results indicate that the boundary between the IMD and the CIL of members of the dispanin/CD225 family is important for the regulation of their trafficking from the Golgi. This domain can be clearly defined in members of the dispanin/CD225 A subfamily as a consensus sequence conserved in vertebrates, and despite sequence dissimilarity, it maintains similar functions in other subfamilies, at least as appreciated in PRRT2 using *de novo* generated as well as PKD-associated genetic mutations.

## Conclusions

In the present study, we have defined a novel domain within the CIL of IFITM3 that regulates egress from the Golgi through the characterization of a novel gain-of-function IFITM3 mutant. This _81_S-x-K-x-R-D_86_ domain is conserved at the amino acid level in vertebrate members of the dispanin/CD225 A subfamily that regroups all IFITM members. In addition, we do show here that, despite no sequence similarity, this domain is nonetheless functionally conserved in at least one member of the B subfamily (PRRT2), raising the possibility of a general functional role across all dispanin/CD225 subfamilies.

Mutations of the Golgi egress domain lead to the retention of IFITM3 in the Golgi, where IFITM3 interferes with v-to t-SNARE vesicles fusion, ultimately driving macroscopic changes in the Golgi apparatus and glycoproteins trafficking defects.

The inhibitory effect of the 85-90 IFITM3 mutant on v-to t-SNARE vesicles membrane fusion is highly reminiscent of the well-described negative effects of IFITM proteins on the fusion between viral and cellular membranes after viral envelopes engage their cellular receptors and in this respect it is not surprising that increased accumulation of IFITM3 at the Golgi affects SNAREs fusion similarly. However, these results stress the importance of a controlled regulation of the exit from the Golgi of IFITM3, and possibly of other members of the family, to avoid deleterious effects on the functionality of the entire apparatus.

While ER-sorting protein domains have been identified^28^, similar domains have not been characterized for the Golgi and our results thus provide an important piece of the puzzle to decorticate how proteins that transit through the Golgi may be sorted out of it.

The Golgi egress domain of IFITM3 can either serve as a docking site for specific co-factors that mediate trafficking within and from the Golgi (for instance Rab proteins), it can play a regulatory role of other IFITM regions, or of course a combination of both.

The _81_S-x-K-x-R-D_86_ domain is specially posited between the IMD and the beginning of the CIL and it is thus in close proximity with a short amphipathic helix important for antiviral functions^29^, with two cysteine residues that can be palmitoylated and thus affect IFITM association to membranes (C71, C72 in human IFITM3; a third cysteine C105 is also present that appears less important in IFITM3 functions^30,31^) and a G-x_3_-G domain that has been shown to be important for IFITMs oligomerization and functions^32^. Furthermore, the K83 residue present within the _81_S-x-K-x-R-D_86_ domain can be itself ubiquitinated 6 and together with residue K104 it has been recently described to mediate PIP3 binding of IFITM3 at the plasma membrane of B cells, in a process that is required for the correct formation of a PI3K signaling platform11. Although, our confocal microscopy data suggest that the K83 IFITM3 mutant may not act as a functional PI3K platform simply because it is retained in the Golgi, the possibility of ubiquitination of the _81_S-x-K-x-R-D_86_, at least in members of the dispanin/CD225 A subfamily, implies the potential prospect of finely tuning Golgi egress according to the dynamic status of ubiquitination of IFITM3, thus adding an additional layer of regulation to this process. Given that the 85-90 IFITM3 mutant is a proficient membrane fusion inhibitor, we believe it unlikely that the _81_S-x-K-x-R-D_86_ domain plays a role in IFITM oligomerization driven by the G-x_3_-G domain, as this would have been expected to lead to a non-functional IFITM3 mutant incapable to inhibit v-to t-SNAREs fusion.

The model we favor is that the _81_S-x-K-x-R-D_86_ domain regulates the access of proximal domains to post translational modifications that in turn affects IFITM trafficking and more specifically access to palmitoylation at proximal cysteines. This model is in agreement with a recent one proposed by the Rothman’s lab in which Golgi exit of certain proteins relies on a gradient of palmitoylation^33^. The model could also explain why this domain plays a conserved functional role in diverse subfamilies, despite no discernable sequence similarity, as is the case for PRRT2. The presence of two cysteines proximal to the region corresponding to the domain that regulates egress from the Golgi in the A subfamily and in PRRT2 is indeed a common feature in all members of the different dispanin/CD225 subfamilies (C276/C278 in PRRT2; C114/116 in TUSC5 as members of the B subfamily; C171/C172 in TMEM90A; C015/C106 in TMEM91 for members of the C subfamily and C152/C153 for PRRT1, member of the D subfamily).

PRRT2 is a neuronal specific protein involved in pre-and post-synaptic neurotransmitter vesicles regulation and has been genetically linked to PKD. While the majority of PKD patients suffer from mutations that result either in absent or severely truncated proteins, a few exhibit non-synonymous changes in the context of the full-length protein. Our results indicate that among them, mutations in the Golgi egress domain of PRRT2 may work by causing relocation of the protein, thus diminishing the concentration of active PRRT2 at its normal functional sites (i.e. the plasma membrane).

Although IFITMs have been predominantly studied in a viral infection context, several reports indicate that these proteins can play more pleiotropic functions: they interfere with trophoblast fusion during placental formation by affecting syncytin-mediated membrane fusion^12,34^; they lead to glucose-related metabolic dysfunctions in mice through unknown mechanisms^35^ and they concur to B cell signaling11. Interestingly, ectopic expression of IFITM proteins has been in some cases reported to decrease the overall levels of viral glycoproteins and even to act more generally on translation, although the underlying mechanism(s) remain unclear^36,37^. In this respect, our results may provide a mechanistical explanation for the decreased levels of glycoproteins, by suggesting that the phenotype can be driven by high levels of IFITM3 accumulation in the Golgi, for instance in conditions of prolonged interferon stimulation, that in turn alter glycoproteins trafficking and stability through this secretory pathway.

More generally, our results highlight the fine line that separates the protective effects of innate defense proteins against viral aggression, from the deleterious side-effects that they may play against the cell itself. The literature provides other examples of this ambivalence for example in the family of apolipoprotein B mRNA editing enzyme, catalytic polypeptide-like 3 proteins (APOBEC3), prototypical anti-retroviral defense proteins. While the main function of APOBEC3 members is to mutate and thus inactivate retroviral genomes thanks to their cytidine deaminase activity, this same enzymatic activity also contributes to the tumorigenic process when it is turned against the cellular genome^38–40^.

IFITMs are potent inhibitors of membrane fusion in the context of viral infection, but since membrane fusion is a common process in the cell, their biology (intended here as the sum of activity, dynamic location, stability etc) must be tightly regulated to avoid deleterious effects on the cell itself. Together with other examples mentioned above and recently described in the literature, our results suggest that dysregulated expression of IFITM molecules and in particular with signals that interfere with its egress from the Golgi may lead to Golgi-related dysfunctions. This may be an as yet unappreciated contribution of IFITM proteins to some pathological conditions for example in cancers or in interferonopathies in which the expression of IFITM proteins appears deregulated.

## Methods

### Cells, plasmids and antibodies

Human embryonic kidney cells (HEK293T, ATCC cat. CRL-3216) were maintained in complete DMEM media with 10% Fetal Calf Serum (Sigma cat. F7524). Untagged WT IFITM3 (gene ID: 10410) and IFITM1 (gene ID: 8519) and corresponding mutants were either previously described (85-90; ^13,41^), or cloned (all others) by standard mutagenesis techniques in the plasmid pQCXIP (Clontech). WT human PRRT2 (gene ID: 112476) was synthesized as N-terminal HA tag fusion protein (Genewiz) in pcDNA3 vector (Thermofisher) and single point mutants were generated on this matrix by standard molecular biology techniques.

The HIV-1 proviral clone NL4-3 and the VSV-G expression constructs have been described elsewhere^42^. The mEmerald-VSV-G (mEm.-VSV-G) coding construct was a gift from Michael Davidson (Addgene, 54307); pcDNA-D1ER coding for an ER targeted enhanced CFP was a gift from Amy Palmer & Roger Tsien (Addgene cat. 36325 ^43^). GS15-CFP and ERS24-YFP (cyanin and yellow fluorescent proteins, respectively) were obtained by Oleg Varlamov from the Oregon National Primate Research Center, USA ^20^. The following primary antibodies were used for WB or confocal microscopy, as indicated. Mouse monoclonals: anti-α-Tubulin, anti-HA and anti-VSV-G (Sigma cat. T5168, cat. H3663 and cat. V5507, respectively), anti-LAMP2 (Santa Cruz Biotechnology, cat. sc-18822), anti-CD63 and anti-GM130 (BD biosciences cat. 556019 and cat. 610823); anti-HIV-1 p24 (obtained through the NIH HIV Reagent Program, Division of AIDS, NIAID, NIH, contributed by Dr. Bruce Chesebro and Kathy Wehrly, cat. ARP-3537). Rabbit polyclonal antibodies: anti-IFITM3 (Proteintech cat. 11714-1-AP), anti-IFITM1 (Proteintech, cat. 60074) and anti-GM130 (abcam cat. Ab52649). The sheep polyclonal antibody anti-HIV-1 Env was obtained through the NIH HIV Reagent Program, Division of AIDS, NIAID, NIH (contributed by Dr. Michael Phelan, cat. ARP-288).

The following secondary antibodies were used for WB: anti-mouse, anti-rabbit and anti-sheep IgG-Peroxidase conjugated (Sigma, cat. A9044 and cat. AP188P and Dako cat. P0163); while the following ones were used for confocal microscopy: donkey anti-rabbit IgG–Alexa Fluor 594 conjugate and donkey anti-mouse IgG–Alexa Fluor 488 conjugate (cat. A-21207 and cat. A-21202; Life Technologies).

### Ectopic DNA transfections, viral production and confocal microscopy analyses

HEK293T cells were directly seeded on 0.01% poly-L-lysine-coated coverslips (Sigma, cat. P4832) and analyzed 24 hours after ectopic DNA transfection (unless otherwise specified, Lipofectamine 3000 cat. L3000008, ThermoFisher, according to the manufacturer’s instructions). Cells were washed three times with PBS 1x, fixed with 4% paraformaldehyde (Euromedex, cat. 15713) for 10 min, quenched with 50 mM NH_4_Cl (Sigma cat. A4514) for 10 min, and permeabilized with PBS–0.5% Triton X-100 (Sigma, cat. X100) for 5 min. After a blocking step in PBS–5% milk, cells were incubated with primary antibodies for 1 hour at room temperature (dilution 1:100), washed and then incubated with fluorescent secondary antibodies (dilution 1:100). A 4’-5-diamidina-2-phenylindole (DAPI)-containing mounting medium was used (ThermoFisher, cat. 62248). Images were acquired using a spectral Zeiss LSM800 confocal microscope and analyzed with Fiji software (version 2.0.0). For 3D-reconstruction, z-stack collections were analyzed with the Imaris 9.2.0 software (Oxford Instruments Group). Colocalisations were quantified using the Pearson overlap coefficient (Fiji software).

IFITMs and PRRT2 coding DNAs were routinely transfected at a concentration of 1 μg per well of a 24 well plate along with 0.2 μg of VSV-G or of mEm.-VSV-G. When specified, increasing doses of GS15 coding DNA were also added (0, 0.2, 0.5 and 1 μg).

Monensin was used at a final concentration of 0.2 mM for 3 hours, prior to analysis (Sigma, cat. M5273).

Production of HIV-1 virions particles was instead carried out by calcium phosphate DNA transfection of the HIV-1 proviral clone NL4-3 (ratio 3 to 1, NL4-3 over IFITM). Virion particles released in the supernatant of transfected cells were harvested forty-eight fours post transfection, syringe-filtered (0.45μm filters, Minisart, cat. 146622) to remove cellular debris and purified by ultracentrifugation over a 25% sucrose cushion (28.000 rpm for 2 hours; Beckman Coulter ultracentrifuge).

In confocal microscopy experiments using the GM130 Golgi marker, IFITM or VSV-G relocalization was quantified for each cell as the ratio between the signal present at the Golgi over the signal present in the cytoplasm (Golgi/cytoplasm ratio). When the GM130 marker was not used, perinuclear accumulation was either scored visually (as in Fig 1, in light of the extremely clear phenotype; binary scoring: perinuclear or non-perinuclear), or quantified by determining the average percentage of accumulation of VSV-G at the left or right part of the nucleus (asymmetric distribution= perinuclear accumulation; symmetric distribution= non-perinuclear accumulation). To bypass artificial measurements due to differences in size between left and right portions of the cell, the avg of the signals measured at each side of the cell were used. This measurement diminishes the absolute values of the protein present at one side, but offers the advantage of making measures independent from cell size differences. Visual scoring in Fig 1 was also confirmed through this analysis.

### Electron microscopy

Samples were fixed in mixture of 4% paraformaldehyde (TAAB Laboratories Equipment Ltd, United Kingdom, cat. F/P001) and 1% glutaraldehyde (Electron Microscopy Science, USA, cat. 16310) in 0.1 M phosphate buffer (pH 7.4) for 24 hours, washed three × 30 min in 0.1 M of phosphate buffer, and post-fixed for one hour with 2% osmium tetroxide (Electron Microscopy Science, USA, cat. 19190) in 0.15 M of phosphate buffer. After washing in 0.1 M of phosphate buffer for 20 min and two x 20 min in distillated H_2_O, samples were dehydrated in a graded series of ethanol solutions (50% ethanol two x 10 min; 70% ethanol three x 15 min and last portion for 14 hours; 90% ethanol three x 20 min; and 100% ethanol three x 20 min). Final dehydration was performed by 100% propylene oxide (PrOx, ThermoFisher (Kandel) GmbH, Germany, Lot X19E013) three x 20 min. Then, samples were incubated in PrOx/EPON epoxy resin (Fluka, Switzerland) mixture in a 3:1 ratio for two hours with closed caps, 16 hours with open caps, and in 100% EPON for 24 hours at room temperature. Samples were replaced in new 100% EPON and incubated at 37 °C for 48 hours and at 60 °C for 48 hours for polymerization. Serial ultra-thin sections (thickness 70 nm) were cut with a “Leica Ultracut UCT” ultramicrotome (Leica Microsysteme GmbH, Wien, Austria), placed on TEM nickel one-slot grids (Agar Scientific, Ltd. United Kingdom, cat. G2500N) coated with Formvar film and stained 20 min with 5% uranyl acetate (Merck, Darmstadt, Germany, cat. 8473) and 5 min Reynolds lead citrate. The sections were then observed at 100 kV with a Jeol 1011 TEM (JEOL, Tokyo, Japan) connected to a Gatan digital camera driven by Digital Micrograph software (GMS 3, Gatan, Pleasanton, CA, USA).

### Fluorescence Resonance Energy Transfer (FRET) microscopy

Analyses were performed on live cells plated on a 35 mm glass-bottom dish and transfected with 2 μg of DNA coding the 85-90 IFITM3 mutant and 0.5 μg each of GS15-CFP and ERS24-YFP to ensure that the majority of cells examined were IFITM3-positive. After replacing media with phenol-red MEM supplemented with 10%FCS, cells were analyzed by confocal microscopy and images were acquired at a temperature of 37°C with a spectral Zeiss LSM710 confocal microscope, equipped with an argon laser and a 63X oil immersion objective. CFP and YFP were excited with a 458 and a 514 nm lasers (emission windows set at 465-505 nm and 525-600 nm, respectively) and images were acquired with a 512×512 resolution, 8 bit line average. Photobleaching of the donor YFP was performed with 4 sequential illuminations of the selected cell (4 frames, line average 2, iterations at 400, scan speed at 8, 514 nm laser AOTF setting at 100%). Four scans were acquired after each bleaching (two before and two after). FRET efficiency was calculated from the ratio of the CFP fluorescence measured post-and pre-YFP bleaching. Analyses were performed on 15–20 individual cells per sample group. The FRET control plasmid pECFP18aaEYFP from addgene.) was used for setup (Addgene cat. 109330, gift from Gabriele Kaminski Schierle) ^44^. Images were analyzed using the ROI manager in Fiji software.

### Glycosidase treatment assays and WB quantification

Cell lysates expressing IFITM3 and VSV-G proteins were collected and split in three aliquots that were either untreated, or treated with Endoglycosidase H (EndoH, NEB, P0702S) or N-glycosilase F (PNGaseF, NEB P0704S), according to the manufacturer’s instructions. Treated lysates were then analyzed by WB, images acquired using Image Lab Touch Software (version 2.0.0.27, Chemidoc Imaging System from Bio-Rad) and bands were quantified by densitometry using the volume tool in the same software.

### Softwares

Electron microscopy: Digital Micrograph software (GMS 3, Gatan, Pleasanton, CA, USA). Confocal microscopy: Fiji software (version 2.0.0), Zen (version 2.3, Zeiss) and Imaris 9.2.0 software (Oxford Instruments Group). WB: Image Lab Touch Software (version 2.0.0.27, Chemidoc Imaging System from Bio-Rad).

Statistics and graphs: Graphpad Prism8 (8.4.3, Graphpad software, LLC).

### Statistical analyses

The statistical analyses used in this study were calculated with the Graphpad Prism8 software: Student t tests (unpaired, two-tailed), one-way Anova tests with either Tukey’s or Dunnett’s multiple comparisons, as indicated in the figure legends.

## Acknowledgements

Thanks to Delphine Muriaux for sharing insights, to Romain Appourchaux for his initial input, to Léa Picard and Lucie Etienne for help with bioinformatic analyses and evolutionary discussions and to Federico Marziali for help with FRET experiments. We are indebted to Oleg Varlamov from the Oregon National Primate Research Center, USA for v-and t-SNARE constructs as well as to the different authors of plasmids retrieved through Addgene cited in the Methods section.

LZ is the recipient of a PhD fellowship of the Chinese Scholarship Council (CSC). Work in the laboratory of AC is supported by grants from the ANRS (AO-2019-1 and AO-2021-1), as well as the ANR (ANR-20-CE15-0025-01). The funders had no role in study design, data collection and analysis, decision to publish, or preparation of the manuscript. We acknowledge the contribution of the microscopy (LYMIC-PLATIM) platform of SFR BioSciences Gerland Lyon Sud (UMS3444/US8).

## Author Contributions

L.Z. designed and performed most of the experiments. R.U. and P.R. performed and analyzed electron microscopy. A.C. supervised the research and wrote the manuscript. All authors commented on the manuscript.

## Data availability

Source data is provided with this paper. There is no restriction on data availability.

## Figure Legends

**Extended data. Figure 1. The 85-90 IFITM3 mutant does not colocalize with CD63, LAMP2 nor with the ER.** HEK293T cells ectopically expressing the indicated IFITM3s, or co-transfected with the ER reporter D1ER were analyzed by confocal microscopy with the indicated markers. Whiskers and boxes plot of the Pearson’s colocalization coefficients (25-75 percentiles with individual cells represented as dots; n=16 to 38 per condition in three independent experiments). A one-way Anova with Tukey’s multiple comparison test was applied for each marker to evaluate differences in localization between WT and 85-90 IFITM3 mutant (non statistical significant differences are represented by white boxes; p <0.0001 by colored ones). Source data is provided in the relevant section.

**Extended data. Figure 2. The broad trafficking inhibitor Monensin affects the egress from the Golgi of WT IFITM3, but not of the 85-90 mutant.** Cells expressing the indicated IFITM3 protein were either treated or untreated with Monensin for 3 hours at 0.2 mM, prior to confocal microscopy analysis. Panels present typical staining, while the graph presents the proportion of IFITM3 in the Golgi as an IFITM3 Golgi/cytoplasmic ratio (AVG= lines and individual values displayed as dots). The lower table presents the result of the indicated statistical analyses: ns, non significant; ** p values ≤ 0.006. Source data is provided in the relevant section.

**Extended data. Figure 3. Complete confocal microscopy analyses of the individual point mutants of the CIL of IFITM3.** This figure presents the entire panel of confocal microscopy analyses (i.e. IFITM3, GM130 and DAPI staining) of the mutants presented in Figure 5, which for lack of space presents only IFITM3 staining panels. Source data is provided in the relevant section of Figure 2.

**Extended data Figure 4. Complete confocal microscopy analyses of the individual point mutants of IFITM1 and PRRT2.** This figure presents the entire panel of confocal microscopy analyses (i.e. PRRT2, IFITM1, GM130 and DAPI staining) of the mutants presented in Figure 6b and 6c, which for lack of space presents only overlays but not separated channels. Source data is provided in the relevant section of Figure 6.

