## Supplementary figures for "A novel domain within the CIL regulates egress of IFITM3 from the Golgi and prevents its deleterious accumulation in this apparatus"

Li et al, Extended Data. Figure 1

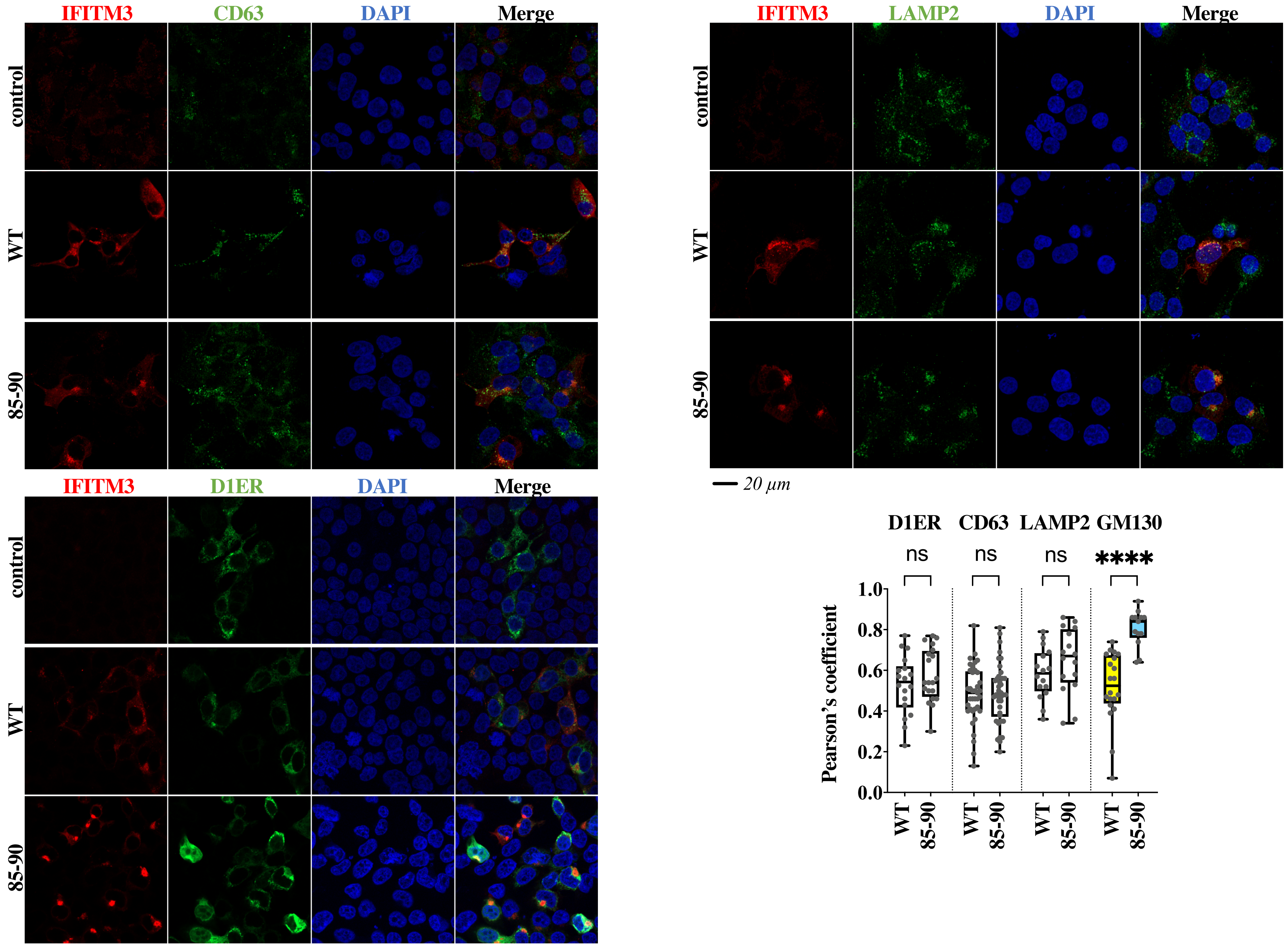

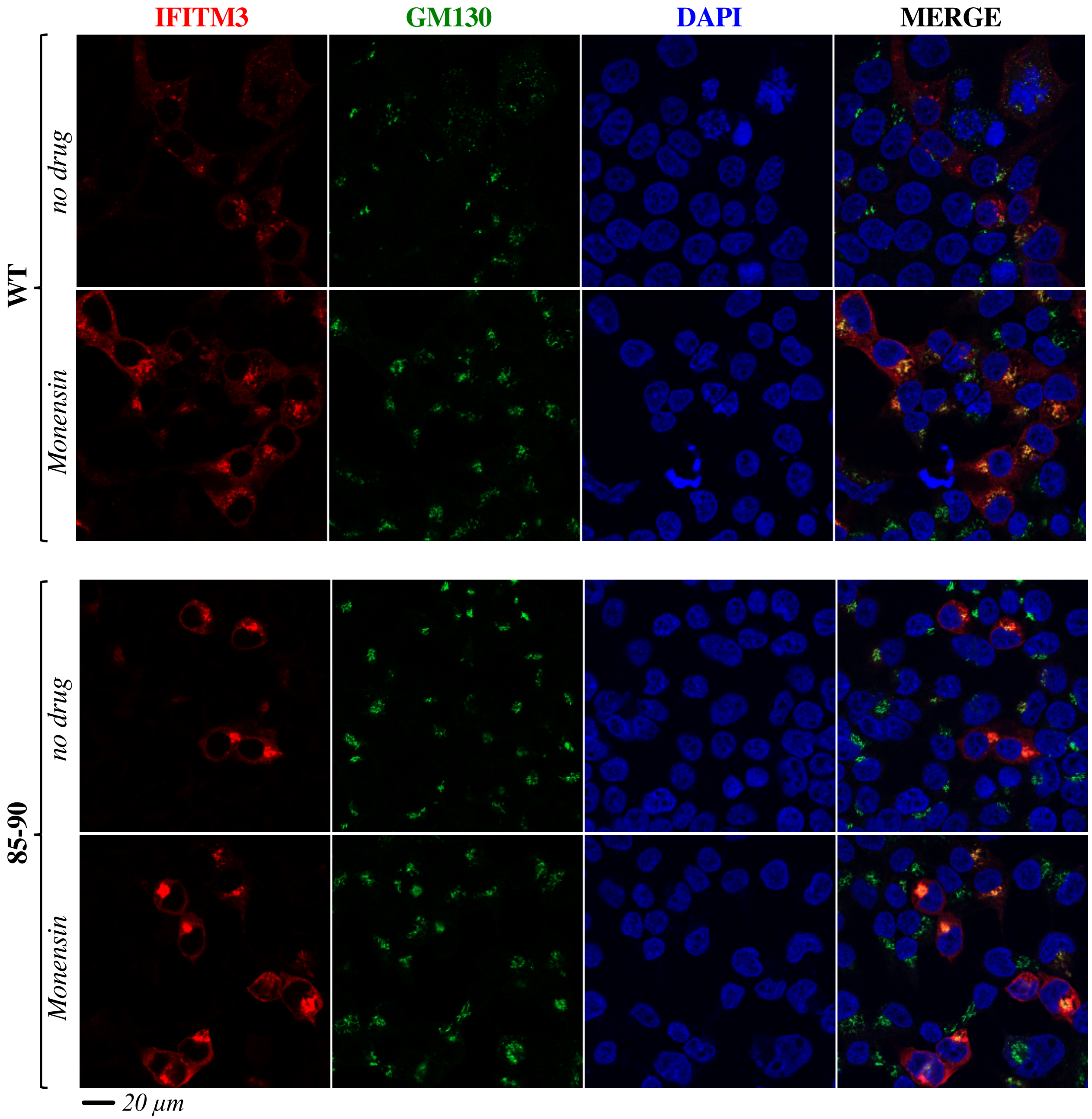


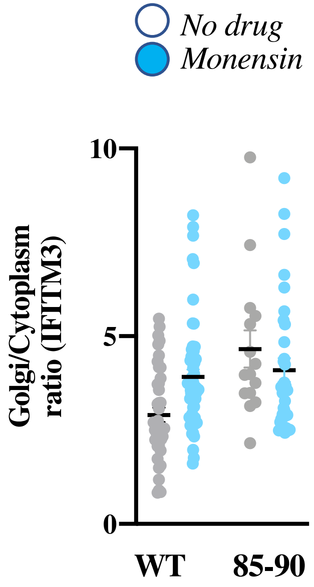


| | WT Monensin | 85-90 Monensin |
|---|---|---|
| WT no drug | ** | ** |
| 85-90 no drug | ns | ns |
| WT Monensin | | ns |
| 85-90 Monensin | ns | |

***one-way Anova, Tukey's multiple comparison test***

*Li et al, Extended Data. Figure 3*

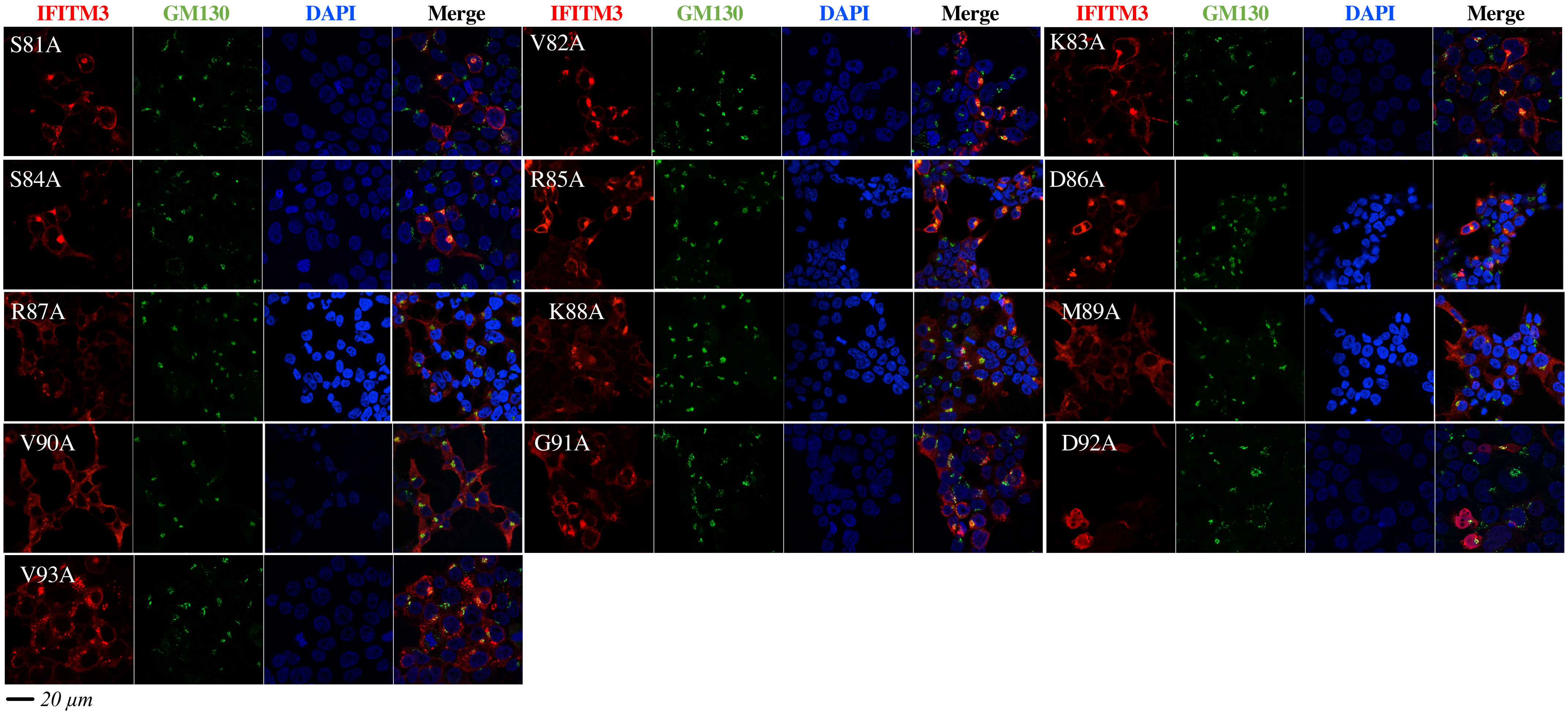

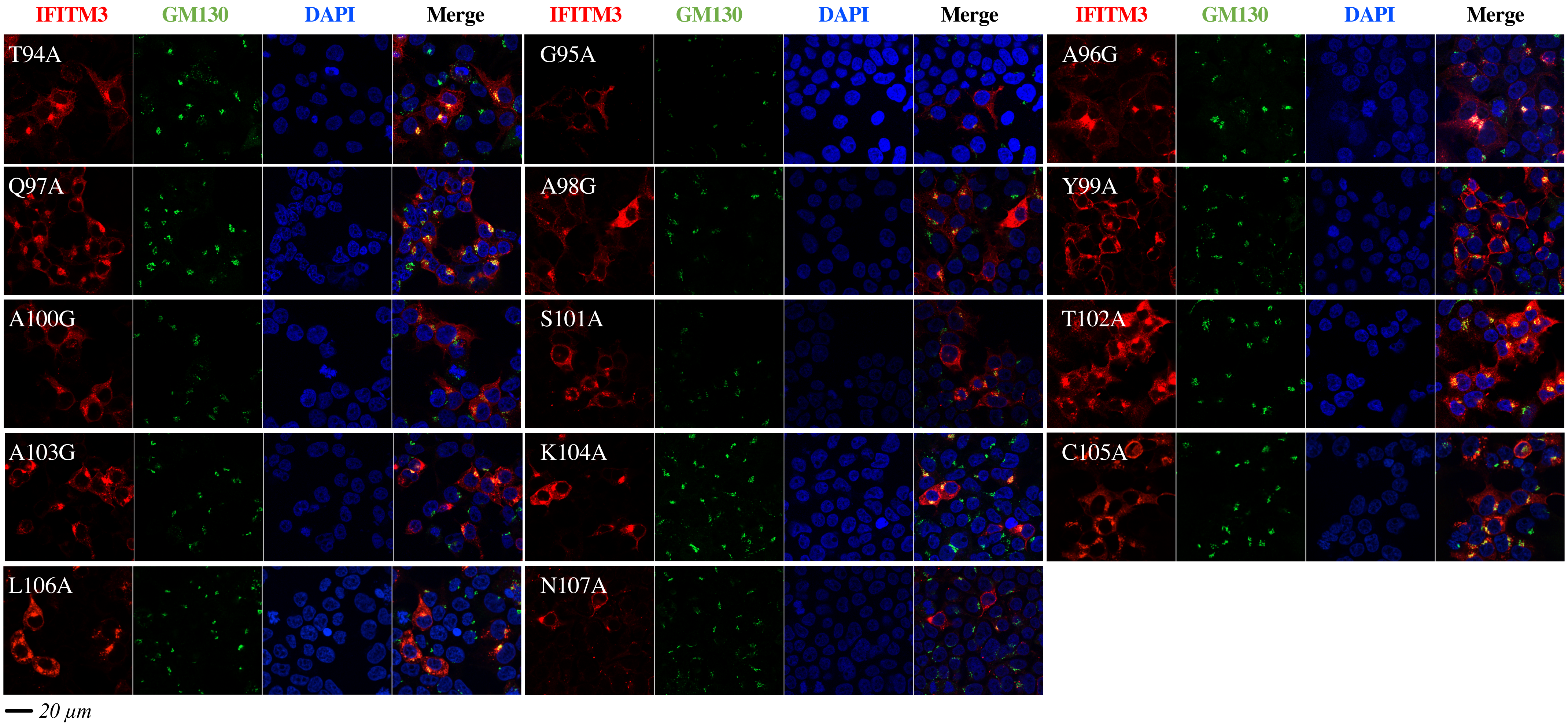
IFITM3
GM130
DAPI
Merge
T94A
G95A
A96G
Q97A
A98G
Y99A
A100G
S101A
T102A
A103G
K104A
C105A
L106A
N107A
20 µm

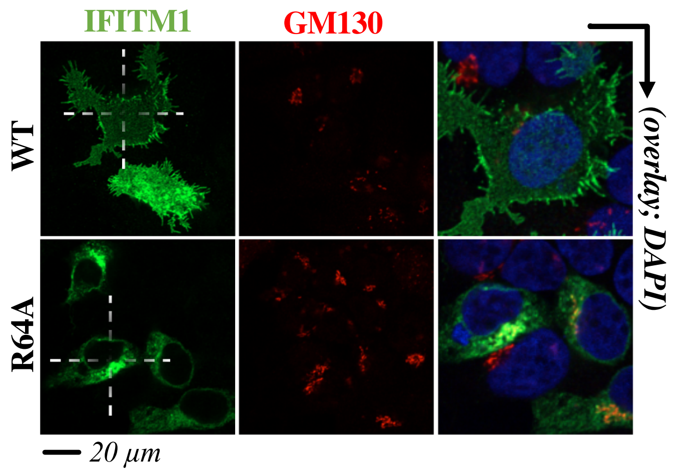
IFITM1
GM130
(overlay; DAPI)
WT
R64A
20 μm

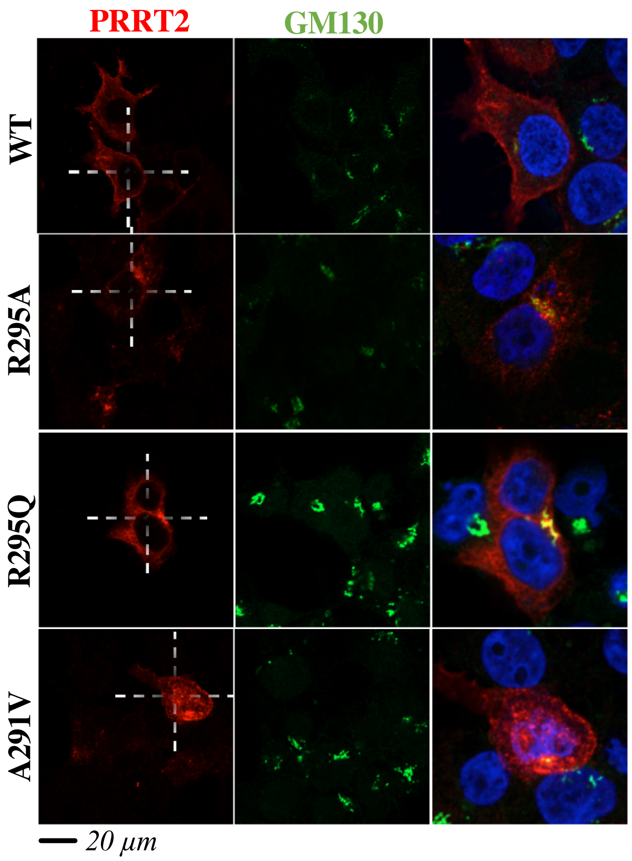
PRRT2
GM130
WT
R295A
R295Q
A291V
20 μm
